# Multiplex serologic testing within a cross-sectional lymphatic filariasis sentinel site survey in coastal Kenya reveals community-level differences in IgG antibody responses to parasitic diseases and vaccines

**DOI:** 10.1101/604181

**Authors:** Sammy M. Njenga, Henry M. Kanyi, Benjamin F. Arnold, Hadley S. Matendechero, Joyce K. Onsongo, Kimberly Y. Won, Jeffrey W. Priest

## Abstract

Accurate, cost-effective measurement of the burden of co-endemic infections would enable public health managers to identify opportunities for implementation of integrated control programs. Dried blood spots (DBS) collected during a cross-sectional lymphatic filariasis sentinel site survey in the Kenyan coastal counties of Lamu, Tana River, Kilifi, Kwale, and Taita-Taveta were used for the integrated detection of serologic IgG antibodies against antigens from several parasitic infections (*Wuchereria bancrofti*, *Schistosoma mansoni*, *Plasmodium* spp, *Ascaris lumbricoides*, and *Strongyloides stercoralis*) as well as markers for immunity to vaccine-preventable diseases (measles, diphtheria, and tetanus) on a multiplex bead assay (MBA) platform. High heterogeneity was observed in antibody responses by pathogen and antigen across the sentinel sites. Antibody seroprevalence against Wb123, Bm14, and Bm33 recombinant filarial antigens were generally higher in Ndau Island (p<0.0001), which also had the highest prevalence of filarial antigenemia compared to other communities. Antibody responses to the *Plasmodium* species antigens CSP and MSP-1_19_ were higher in Kilifi and Kwale counties, with Jaribuni community showing higher overall mean seroprevalence (p<0.0001). Kimorigo community in Taita-Taveta County was the only area where antibody responses against *Schistosoma mansoni* Sm25 recombinant antigen were detected. Seroprevalence rates to *Strongyloides* antigen NIE ranged between 3% and 26%, and there was high heterogeneity in immune responses against an *Ascaris* antigen among the study communities. Differences were observed between communities in terms of seroprevalence to vaccine-preventable diseases. Seroprotection to tetanus was lower in all 3 communities in Kwale County compared to the rest of the communities. This study has demonstrated that the MBA platform holds promise for rapid integrated monitoring of trends of infections of public health importance in endemic areas, and assessing the effectiveness of control and elimination programs.

**Author Summary:** Establishment of successful private-public partnerships in the recent past has led to an increase in resources available for control and elimination of malaria and Neglected Tropical Diseases (NTDs). Implementation of control and elimination programs and their subsequent monitoring and evaluation would be greatly facilitated by development of new tools and strategies for rapid identification of areas of transmission so that interventions could be prioritized to regions where they were most needed. Since development of antibody responses in a host depend on exposure to an infectious agent, assessment of such serologic markers provides a sensitive way to measure differences between populations in pathogen exposure. Our study applied a state-of-the-art multiplex bead assay platform to perform integrated measurement of antibody responses to multiple parasitic diseases and immunizing antigens for vaccine-preventable diseases (VPDs) in ten lymphatic filariasis sentinel sites across the Kenyan coastal region. A community-level analysis of age-specific and overall mean seroprevalence fit using a flexible model ensemble provided an improved understanding about the distributions of the various parasitic infections and seroprotection to VPDs. This study provides an important proof of concept for how we could dramatically increase the value of existing surveillance activities using small volumes of blood collected on filter paper and analyzed using a single multiplex laboratory assay and novel data analysis techniques.

## Introduction

Persons living in tropical and subtropical areas are often faced with enormous health challenges resulting from the co-endemicity of HIV/AIDS, tuberculosis, and malaria. In addition, several other infectious diseases found in sub-Saharan Africa including some Neglected Tropical Diseases (NTDs) are common, particularly among the poor [1–3]. Past studies in the region have identified subgroups who are polyparasitized with soil-transmitted helminth (STH) infections, filarial parasites, and malaria [4–6]. Lymphatic filariasis (LF) caused by *Wuchereria bancrofti* is principally confined to the coastal region of Kenya where ecological factors are suitable for its transmission [7]; the disease co-occurs with other infectious diseases such as STH infections, schistosomiasis, lower respiratory infections, and malaria [8–10].

In the past, lack of resources and competing health priorities in sub-Saharan Africa have led to insufficient commitments to control NTDs. More recently, implementation of successful public-private partnerships (PPPs) for health have availed resources for control and/or elimination of NTDs as public health problems. In 2000 the World Health Organization (WHO) Global Programme to Eliminate Lymphatic Filariasis (GPELF), launched in response to World Health Assembly resolution WHA50.29, urged Member States to initiate activities to eliminate LF as a public health problem, a goal subsequently targeted for 2020 [11]. Community-wide mass drug administration (MDA) of antifilarial drugs for 4-6 years is recommended for LF elimination, and modeling studies have estimated adequate treatment coverage to be at least 65% of total population in endemic areas [12,13]. Substantial progress has been made towards elimination of LF, with Togo being the first country in sub-Saharan Africa to be recognized by WHO for eliminating the disease as a public health problem [14,15]. The Kenyan Ministry of Health launched an LF elimination program in 2002, but the program did not sustain MDA campaigns annually as per GPELF recommendations [16,17]. In 2015, the Ministry of Health successfully appealed to World Health Organization Regional Office for Africa (WHO-AFRO) and other partners for support to re-establish annual MDA campaigns. Subsequently, the WHO Country Office selected the Eastern and Southern Africa Centre of International Parasitic Control (ESACIPAC), which is part of the Kenya Medical Research Institute (KEMRI), to conduct a comprehensive epidemiological assessment of LF infection before re-starting MDA.

Antibody levels can provide valuable information about exposure to infections and can be helpful for characterizing pathogen transmission dynamics to help identify where interventions are needed the most. As some parasite antigens are known to elicit an immunoglobulin G (IgG) response that can be detected for a long period of time, serological analysis of young children provide an estimate of more recent exposure [18,19]. A state-of-art multiplex bead assay (MBA) serological platform that enables simultaneous detection of antibodies against multiple antigens using a small volume of blood sample dried on filter paper [10 μL dried blood spots (DBS)] has been developed as a tool for integrated biomarker surveys [20–22]. The MBA has successfully been used to simultaneously measure antibody responses to multiple parasitic diseases of public health importance as part of a vaccine-preventable disease serological survey in Cambodia [23]. The platform has also been used to simultaneously assess IgG responses to a panel of malaria antigens [24,25]. In the current study, the MBA platform was used for multiplex serosurveillance of diseases of public health importance by testing for antibodies against LF and several other parasitic diseases (malaria, schistosomiasis, ascariasis, strongyloidiasis) as well as seroprevalence to selected vaccine-preventable diseases (measles, diphtheria, and tetanus).

## Methods

### Study design and samples

The DBS samples used in this study were collected during a cross-sectional LF survey conducted in October 2015 in ten sentinel sites located across the coastal region in Taita-Taveta, Kwale, Kilifi, Tana River and Lamu counties as previously described [17]. Briefly, 300 persons aged 2 years or more in each sentinel site were targeted for the LF survey as recommended in the WHO guidelines [11]. The characteristics of the study participants are described in Njenga et al. [17]. The middle finger of consenting individuals was cleaned using a cotton ball soaked in 70% isopropyl alcohol. After drying, the tip of the finger was pricked using a sterile lancet and blood was collected into capillary tubes for detection of circulating filarial antigen (CFA) by immunochromatographic card test (ICT) and onto filter paper for preparation of dried blood spots (6 spots of 10 μl each; Tropbio Pty Ltd, Queensland, Australia) which were used for the MBA.

### Ethical considerations

The study received ethical approval from Kenya Medical Research Institute (KEMRI) Scientific and Ethics Review Unit (SSC No. 3018). In the study villages, chiefs and assistant chiefs arranged for community mobilization meetings during which the purpose of the survey and procedures to be followed were explained. Written informed consent was obtained from every individual who agreed to participate in this study; parents or legal guardians provided signed informed consent forms on behalf of children under 18 years of age. All of the community acquired samples were assayed in the KEMRI-ESACIPAC laboratory in Nairobi, Kenya.

### Recombinant antigens and coupling to microsphere beads

Recombinant *Schistosoma mansoni* glutathione-*S*-transferase (GST) protein was expressed from pGEX 4T-2 plasmid (GE Healthcare, Piscataway, NJ) and purified as previously described [26]. GST fusion proteins that included protein sequences from *Brugia malayi* [Bm33; [21] and Bm14; [19]], *Strongyloides stercoralis* [NIE; [27]], and *Plasmodium falciparum* 3D7 strain [MSP1_19_; [28]] were expressed and purified as previously described. A *W. bancrofti* Wb123-GST fusion protein was a kind gift from T. Nutman (NIH, Bethesda, MD). These proteins were coupled to SeroMap beads (Luminex Corp., Austin TX) using the protein quantities and buffer conditions previously described [23]. *S. mansoni* native soluble egg antigen (SEA) was a kind gift of E. Secor (CDC, Atlanta, GA), and recombinant *S. mansoni* Sm25 antigen was expressed using the Baculovirus system previously described [29]. Both proteins were coupled to SeroMap beads using the protein quantities and buffer conditions previously described [29].

Tetanus toxoid (Massachusetts Biological Laboratories, Boston, MA), diphtheria toxoid from *Corynebacterium diphtheriae* (List Biological Laboratories, Campbell, CA), and recombinant measles nucleoprotein (MV-N, Meridian Life Sciences, Memphis, TN) [30] were purchased from commercial sources. Tetanus toxoid was coupled to SeroMap beads as previously described [31]. Diphtheria toxoid was coupled in buffer containing 50 mM 2-(N-morpholinoethanesulfonic acid (MES) at pH 5.0 with 0.85% NaCl at a concentration of 60 μg of protein per 1.25 × 10^7^ beads in 1 ml final volume. In order to decrease background reactivity, measles MV-N was purified by chromatography on a MonoQ HR 5/5 strong anion exchange column (GE Healthcare, Piscataway, NJ) prior to use. Protein (0.75 mg) was loaded onto the column at a flow rate of 1 ml/ min and washed with 4 ml of 25 mM Tris buffer at pH 8.0. This was followed by a 10 ml linear gradient to 0.25 M NaCl in Tris buffer, then by a 5 ml linear gradient to 1 M NaCl in Tris buffer. The majority of antibody-reactive MV-N eluted in the high salt fractions between 0.4 and 0.7 M NaCl. These fractions were pooled, concentrated using a Centricon-30 centifugal filter device (Millipore Corporation, Bedford, MA), and exchanged into buffer containing 10 mM sodium phosphate with 0.85% NaCl at pH 7.2 (PBS). Approximately 115 μg of protein was recovered (BCA micro assay, Pierce, Rockford, IL). MonoQ purified MV-N was coupled in buffer containing 50 mM MES at pH 5.0 with 0.85% NaCl at a concentration of 6 μg of protein per 1.25 × 10^7^ beads in 1 ml final volume.

Purified native hemoglobin (Hb) from *Ascaris suum* worms was a kind gift from P. Geldhof (Ghent University, Belgium) [32,33]. This antigen was coupled to 1.25 × 10^7^ SeroMap beads in PBS buffer (pH 7.2) at a concentration of 120 μg/ ml.

Cloning of the *P. malariae* MSP1_19_ coding sequence from China I parasite strain is described elsewhere (Priest et al., in preparation). This antigen was coupled to 1.25 × 10^7^ SeroMap beads in 50 mM MES buffer at pH 5.0 with 0.85% NaCl at a concentration of 30 μg/ ml. The glutaraldehyde protocol of Benitez et al. [34] was used to cross-link a synthetic 20 amino acid peptide [(NANP)_5_-amide] corresponding to the carboxy-terminal repeat of the *P. falciparum* circumsporozoite protein (PfCSP) [35,36] to purified GST protein. Bead coupling conditions for this antigen were identical to those described above for the *P. malariae* MSP1_19_ protein.

### Multiplex bead assay

One bloodspot from each person, corresponding to about 10 μl of whole blood, was eluted overnight at 4°C with 200 microliters of PBS containing 0.05% Tween-20 and 0.05% sodium azide (1:40 serum dilution assuming a 50% hematocrit). A further dilution of 50 microliters of eluate into 450 μl of PBS containing 0.5% casein, 0.3% Tween 20, 0.02% sodium azide, 0.5% polyvinyl alcohol (PVA), and 0.8% polyvinylpyrrolidone (PVP) (designated as PBN1) with 3 micrograms/ml *Escherichia coli* extract was made for a final serum dilution of 1:400. Serum dilutions were centrifuged at maximum speed to pellet the *E. coli* extract particulates immediately before use. Bloodspot dilutions were assayed in duplicate with antigen-coupled microsphere beads using a BioPlex 200 system platform (Bio-Rad, Hercules, CA) as previously described [21,23,25]. The average of the median fluorescent intensity values from the duplicate wells *minus* the background fluorescence from the buffer-only blank was reported as the “median fluorescence intensity *minus* background” (MFI-bg). Samples having a coefficient of variation of >15% for ≥2 positive responses between the duplicate wells were repeated.

### Cutoff determinations

WHO International Standard reference sera for tetanus (TE-3; 120 IU/ml) and diphtheria (10/262; 2 IU/ml) purchased from the National Institute for Biological Standards and Control (NIBSC) (Potters Bar, Hertfordshire, United Kingdom) were used to identify MFI-bg cutoff values corresponding to immunoprotection. A tetanus TE-3 value of 10 mIU/ml [37,38] corresponded to a tetanus toxoid MBA response of 118 MFI-bg units. A diphtheria toxoid MBA response of 4393 MFI-bg units corresponded to the 0.1 IU/ml threshold for complete protection [39], and an MBA response of 183 MFI-bg units corresponded to the 0.01 IU/ml threshold for partial protection. Others have shown good concordance between the ‘gold standard’ assays for tetanus and diphtheria and assays using the multiplex bead format [31,40]. Although a WHO reference standard is available for the quantitation of measles virus-neutralizing antibody responses using the whole virus Plaque Reduction Neutralization Test (PRNT) (NIBSC 97/648; 3 IU/ml), the standard has not been calibrated for use in ELISA format assays [41], and our MBA only detects IgG antibodies to the measles MV-N protein. In independent work using the specific bead set from this study Coughlin et al. (in preparation) determined that an ROC-optimized MFI-bg cutoff value of 178 MFI-bg units provided good sensitivity and specificity compared to the ‘gold standard’ PRNT.

MBA cutoff estimates for the *S. stercoralis* NIE assay and for the three LF antigens (Bm33, Bm14, and Wb123) were assigned using a panel of 94 presumed negative sera donated by anonymous adult US citizens with no history of foreign travel. Test values greater than the mean *plus* three standards deviations of the presumed negative sample values were considered to be positive. For the *P. malariae* and *P. falciparum* MSP1_19_ assays, log transformed data were used for the mean *plus* three standard deviation calculation, and the panel used for the *P. falciparum* cutoff included only 65 of the original 94 US adult volunteers. An ROC curve using sera from 41 stool-confirmed, anonymous ascariasis patients, 65 of the adult US citizen volunteers and sera from 45 anonymous US children was used to identify the cutoff for the *Ascaris* Hb MBA. All of the parasitic disease cutoff values were adjusted to account for differences between the instrument used for cutoff determination at the CDC in Atlanta, GA, and the instrument used to assay the Kenyan sample set at KEMRI in Nairobi, Kenya. Two-fold serial dilutions of the same strong positive sera were assayed on both instruments to generate standard curves for cutoff value adjustment.

*S. mansoni* SEA and Sm25 coupled beads were used in an earlier study, and the adjusted, ROC-assigned cutoff values have been reported elsewhere (965 and 38 MFI-bg units, respectively) [29].

We also estimated seropositivity cutoff points for malaria, LF, and helminth antibody responses using the mean plus three standard deviations of a seronegative distribution estimated from the study measurements using finite Gaussian mixture models with two components [42].

### Statistical analysis

Mean antibody levels (MFI-bg) were analyzed on the log_10_ scale due to skewness in their distribution. We estimated age-dependent mean antibody levels and seroprevalence for each study community using cross-validated, ensemble machine learning, with a library that included the simple mean, linear models, locally weighted regression (loess), and smoothing splines with 2 to 10 degrees of freedom, selected using 10-fold cross-validation [43]. We estimated age-adjusted geometric mean antibody levels and seroprevalence for each community using targeted maximum likelihood estimation with influence curve-based standard errors [43]. In cases where seroprevalence approached zero, we estimated exact binomial confidence intervals. Analyses were conducted using R version 3.3.1, and full replication files (data, scripts) are available through the Open Science Framework (https://osf.io/taknp).

## Results

Antibody measurements were obtained from 2,837 individuals (range 271 – 297 per community) (S1 Fig). Antibody distributions varied by pathogen and antigen, and overall there was good concordance between seropositivity cutoff values for malaria, LF and helminth antibody responses derived through ROC curve analysis or mean *plus* 3 standard deviation calculations and those derived by Gaussian mixture model analysis (Figure 1). We therefore relied on cutoff values derived from the Gaussian mixture model antibody responses for comparability to future studies that may not have access to positive and negative control specimens. Age-dependent patterns and community-level estimates of mean antibody levels and seroprevalence were highly consistent (S2 Fig, S3 Fig, S4 Fig, S5 Fig, S6 Fig), so we report results based on mean antibody levels in supporting information.

**Figure 1:**
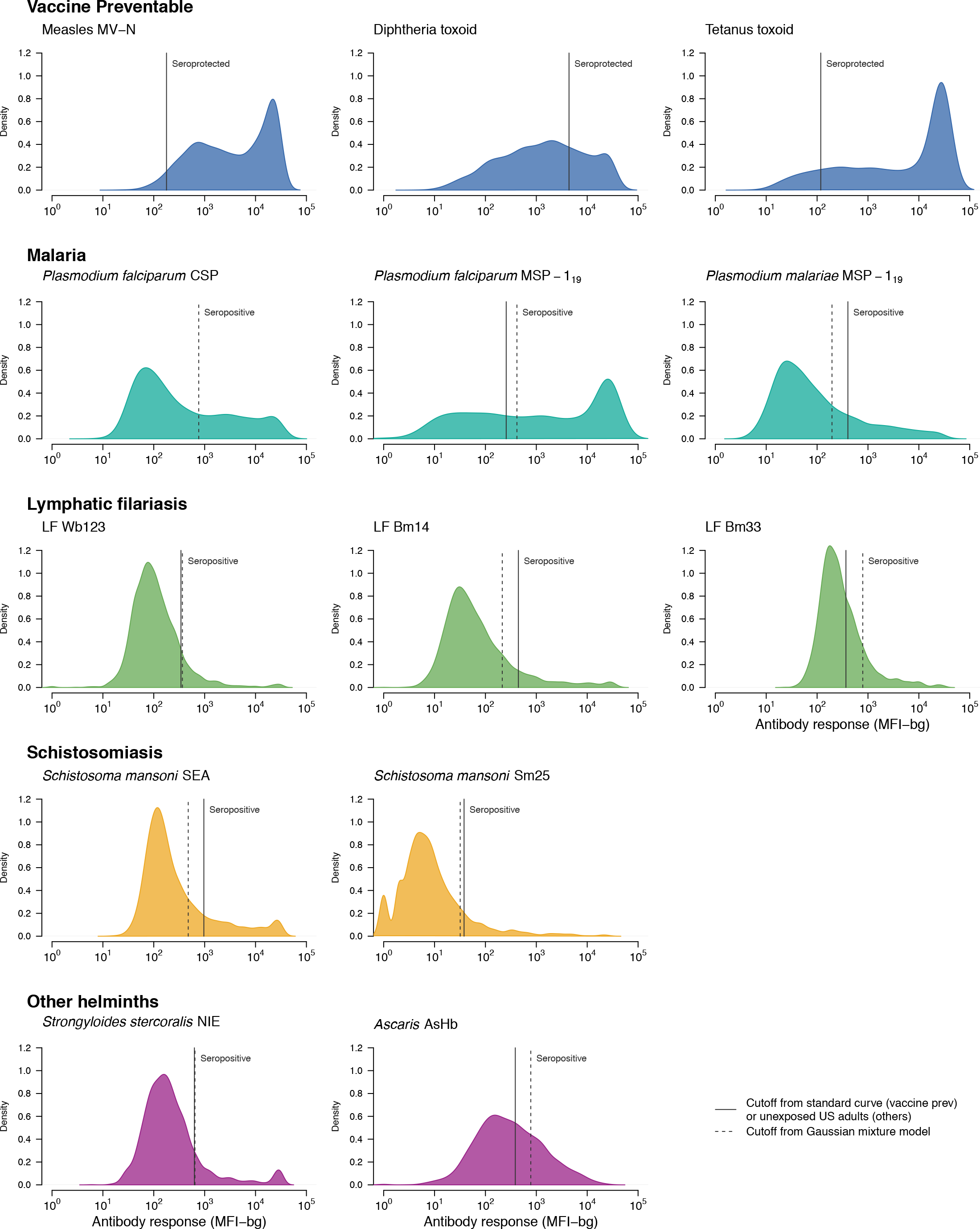
Distribution of quantitative antibody levels measured in 10 communities in Kenya’s coastal region, 2015. Antibody response measured in multiplex using median fluorescence units minus background (MFI-bg) on a BioRad Bio-Plex platform. Seroprotection cutoff points for measles, diphtheria, and tetanus estimated using a standard curve from WHO reference standards. Seropositive cut points for other antigens estimated using negative control serum samples (solid) and finite Gaussian mixture models (dashed). There was no negative control cutoff point determined for the *P. falciparum* CSP antigen. Table S1 includes cutoff values. The script that created this figure is here: https://osf.io/d9jr.

### Antifilarial antibody measurements

Individuals who tested positive for LF infection by ICT had higher mean levels of antibody responses against the 3 recombinant filarial antigens (S7 Fig). Antibody seroprevalence against all 3 recombinant filarial antigens were significantly higher in Ndau Island compared to other communities and the difference in seroprevalence in Ndau compared to other communities was greater among persons less than 30 years old (Figure 2). Antifilarial antibody responses against Bm14 antigen continued to increase with age in all communities. For Wb123, seroprevalence gradually increased with age in Ndau and increased from around the age of 30 - 35 years in Mwadimu community. Compared to the other communities, Jaribuni had slightly elevated mean antibody responses against Wb123 and Bm33 antigens (p<0.0001), but not for Bm14 antigen (p=0.08). Amongst the youngest children, quantitative antibody levels differentiated communities more clearly than seroprevalence owing to high variability in seroprevalence estimates from the small sample sizes in the youngest age strata (S8 Fig). Elevated antibody levels among young children in Ndau, and possibly Jaribuni, were consistent with ongoing LF transmission.

**Figure 2:**
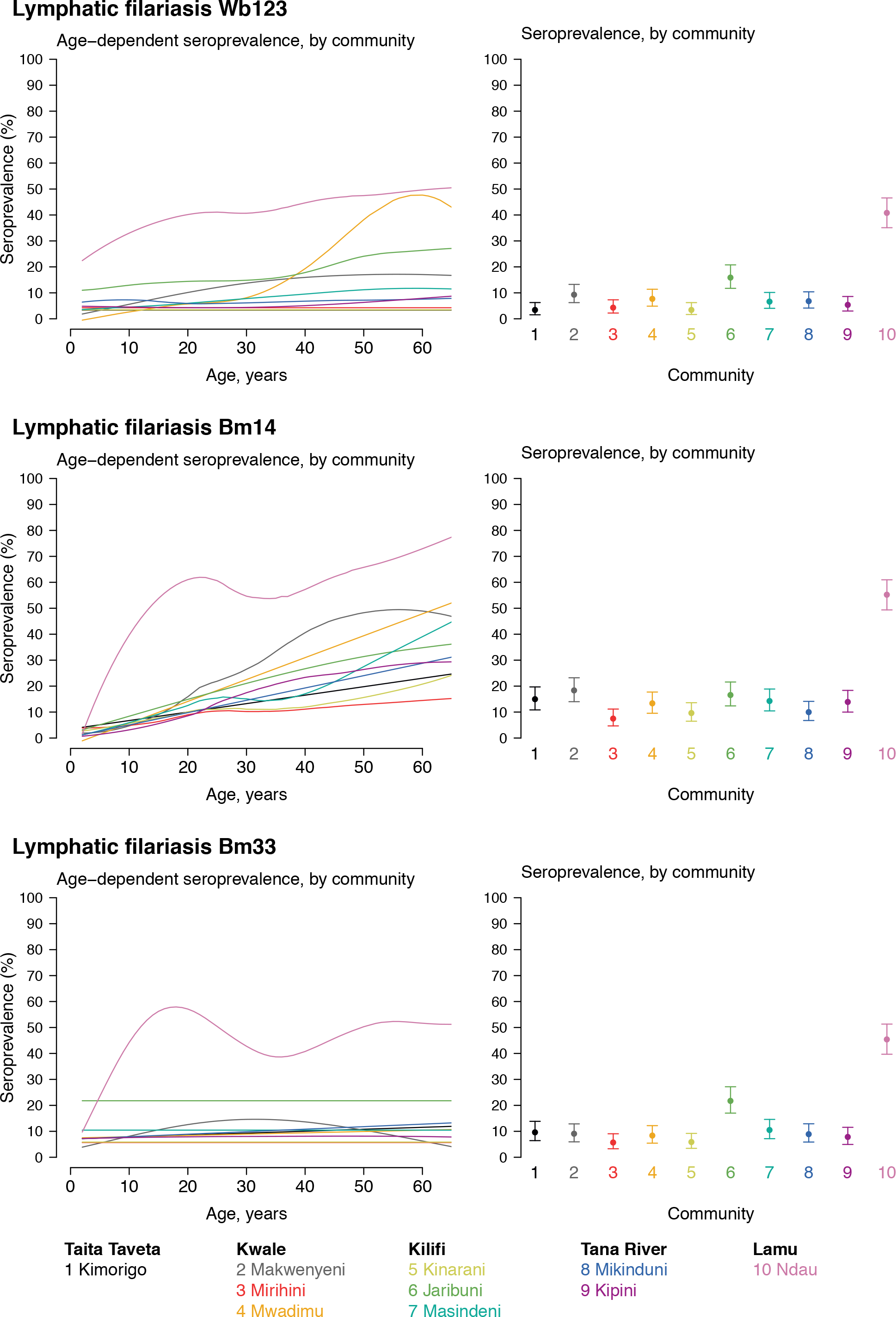
Lymphatic filariasis antibody age-dependent seroprevalence and overall means, stratified by community in Kenya’s coastal region, 2015. Community-level mean seroprevalence is age-adjusted and error bars represent 95% confidence intervals. Figure S2 is an extended version of this figure that also includes quantitative antibody levels. The script that created this figure is here: https://osf.io/5zkxw.

### Antibody responses to other parasite antigens

Antibody responses to the *P. falciparum* CSP and MSP-1_19_ antigens and to the *P. malariae* MSP-1_19_ antigen increased with age in communities in Kilifi and Kwale counties, with higher seroprevalence in Jaribuni community compared to other communities in Kilifi (p<0.0001, Figure 3). Mean antibody responses against *P. malariae* MSP-1_19_ antigen also increased with age and were highest in Jaribuni (p<0.0001), but very low in Ndau Island and Kipini communities (p<0.0001 for difference with other communities).

**Figure 3:**
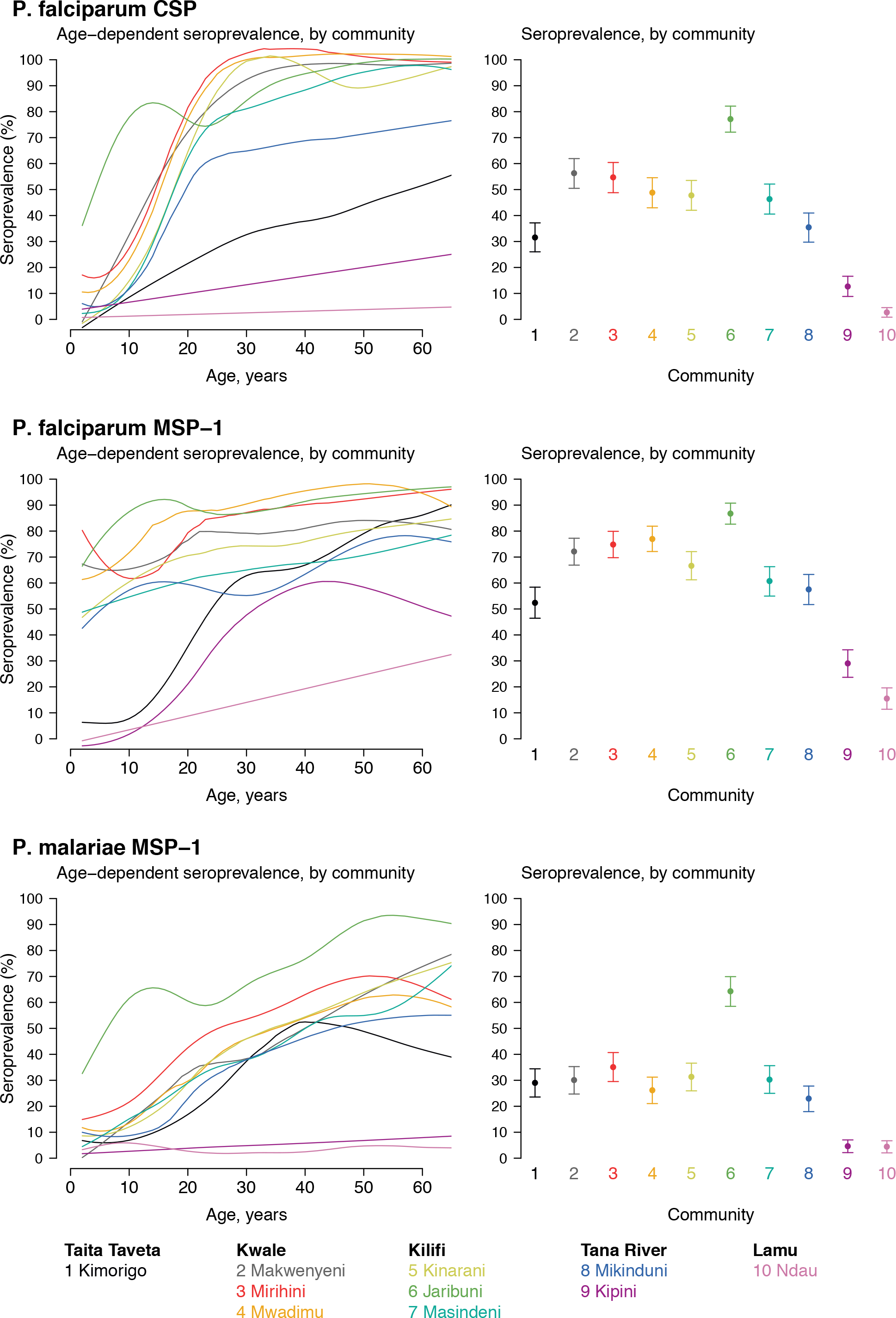
Malarial antibody age-dependent seroprevalence and overall means, stratified by community in Kenya’s coastal region, 2015. Community-level mean seroprevalence is age-adjusted and error bars represent 95% confidence intervals. Figure S3 is an extended version of this figure that also includes quantitative antibody levels. The script that created this figure is here: https://osf.io/kzfd3.

Antibody responses against *Schistosoma mansoni* Sm25 recombinant antigen were primarily detected in Kimorigo community, and the seroprevalence increased gradually with age, reaching a peak at around 25 years of age (Figure 4). However, although antibody responses to *S. mansoni* SEA antigen also increased with age in Kimorigo community and mean seroprevalance was higher, there were some responses against this antigen in many other communities.

**Figure 4:**
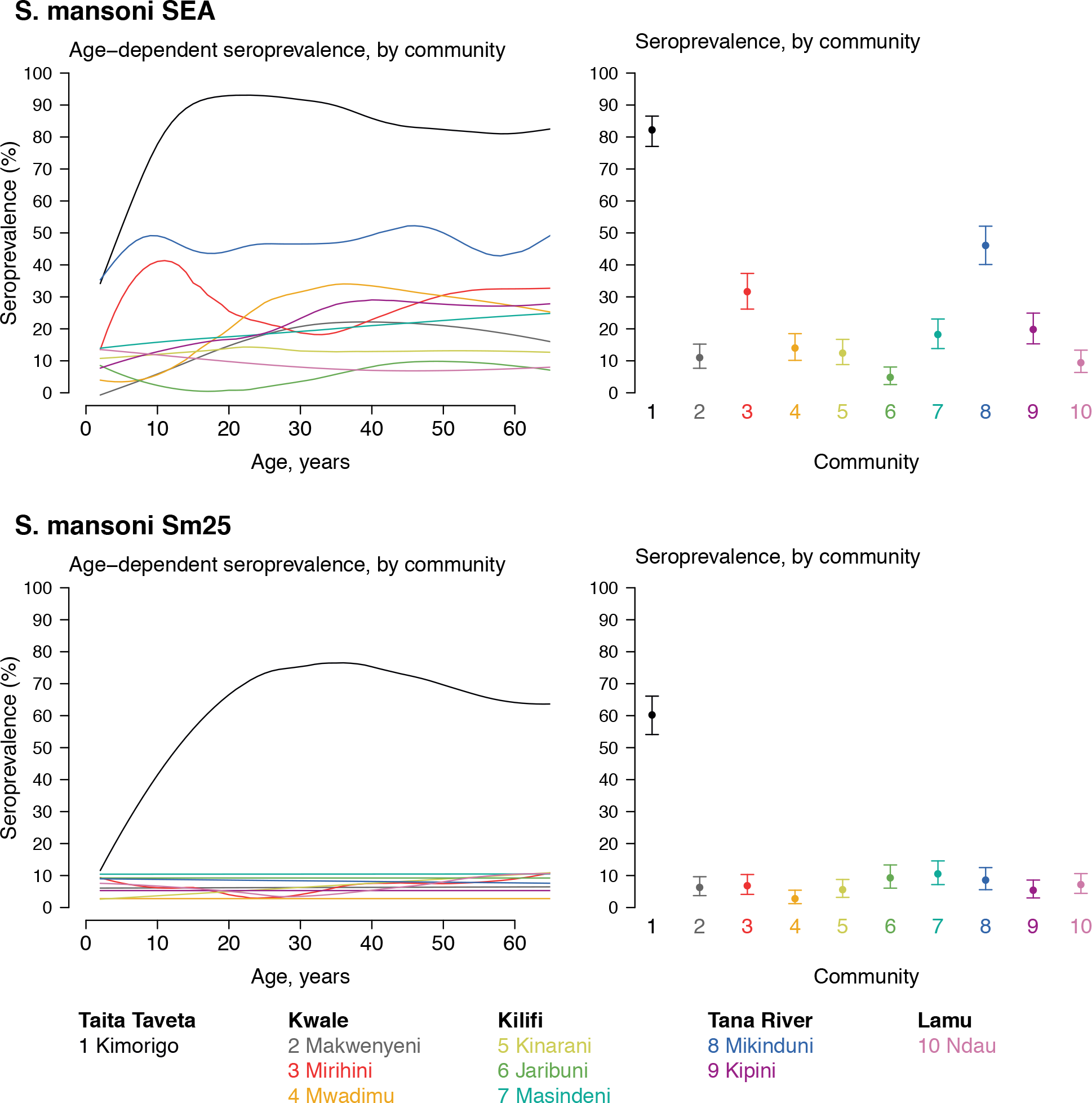
Schistosomiasis antibody age-dependent seroprevalence and overall means, stratified by community in Kenya’s coastal region, 2015. Community-level mean seroprevalence is age-adjusted and error bars represent 95% confidence intervals. Figure S4 is an extended version of this figure that also includes quantitative antibody levels. The script that created this figure is here: https://osf.io/tpcg7.

Steady increases in *S. stercoralis* NIE seroprevalence with age were observed and community level mean seroprevalence ranged between 3% and 26% (Figure 5). There was heterogeneity in age-dependent *Ascaris* Hb seroprevalence patterns across communities, with seroprevalence increasing with age in some communities and decreasing with age in others (Figure 5).

**Figure 5:**
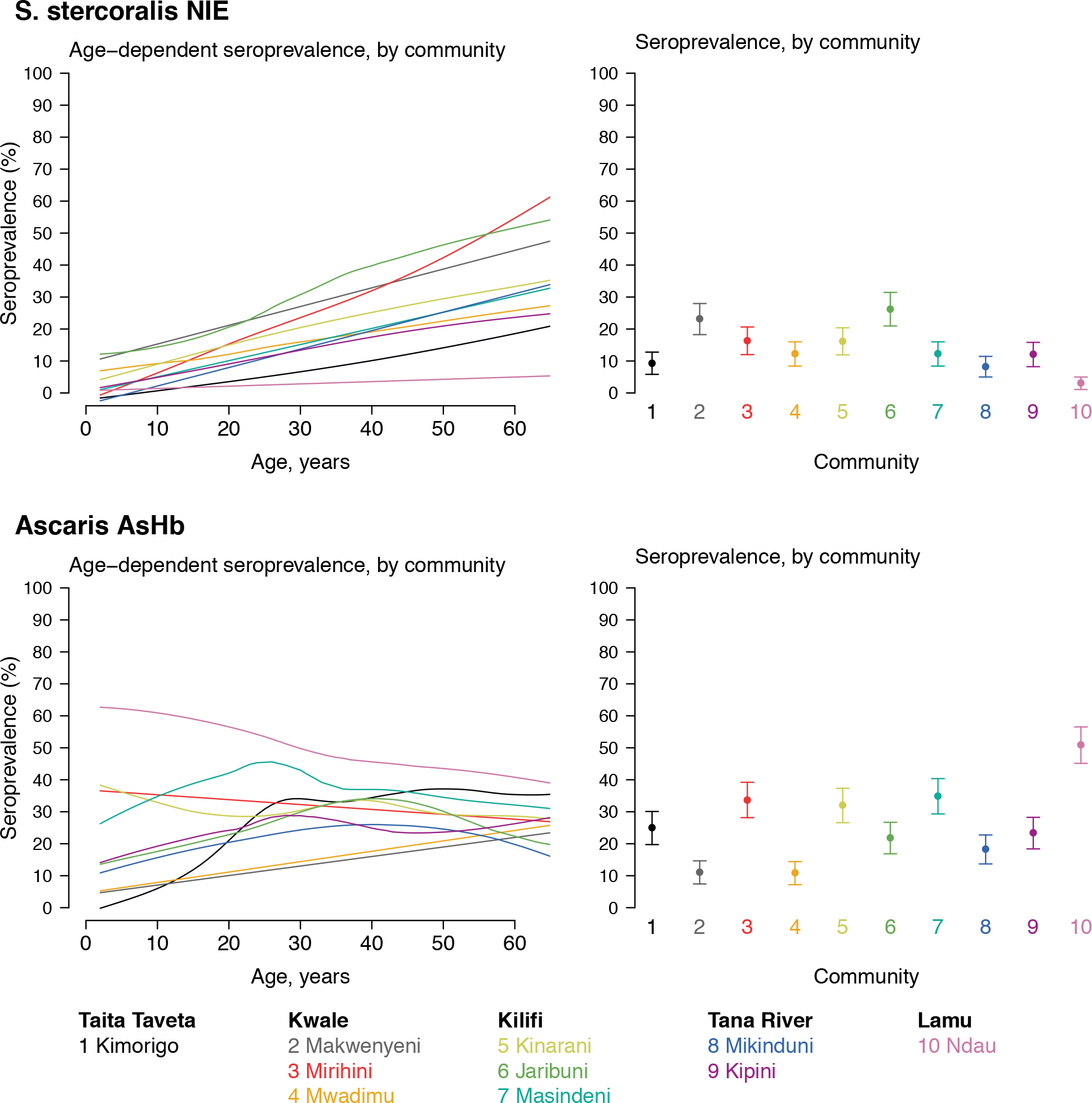
Age-dependent seroprevalence and overall mean for antibodies to S. stercoralis and *A. lumbricoides*, stratified by community in Kenya’s coastal region, 2015. Community-level mean seroprevalence is age-adjusted and error bars represent 95% confidence intervals. Figure S5 is an extended version of this figure that also includes quantitative antibody levels. The script that created this figure is here: https://osf.io/j7ux3.

### Immune responses to vaccine preventable diseases

Immune response against measles MV-N antigen increased with age, but two communities in Kwale County (Mirihini and Mwadimu) had <90% seroprotection (Figure 6). Immune responses to diphtheria toxoid were relatively higher among children, but waned slightly around the ages of 30-40 years before increasing slightly. Generally, diphtheria seroprotection ranged between 22-44% across communities, and partial protection (defined as responses of 0.01-0.099 IU/ml) ranged between 70-88% across communities. Immune responses against tetanus toxoid decreased by age in all communities until around 15 years when the levels increased again. Tetanus seroprotection was lower in all 3 communities in Kwale County.

**Figure 6:**
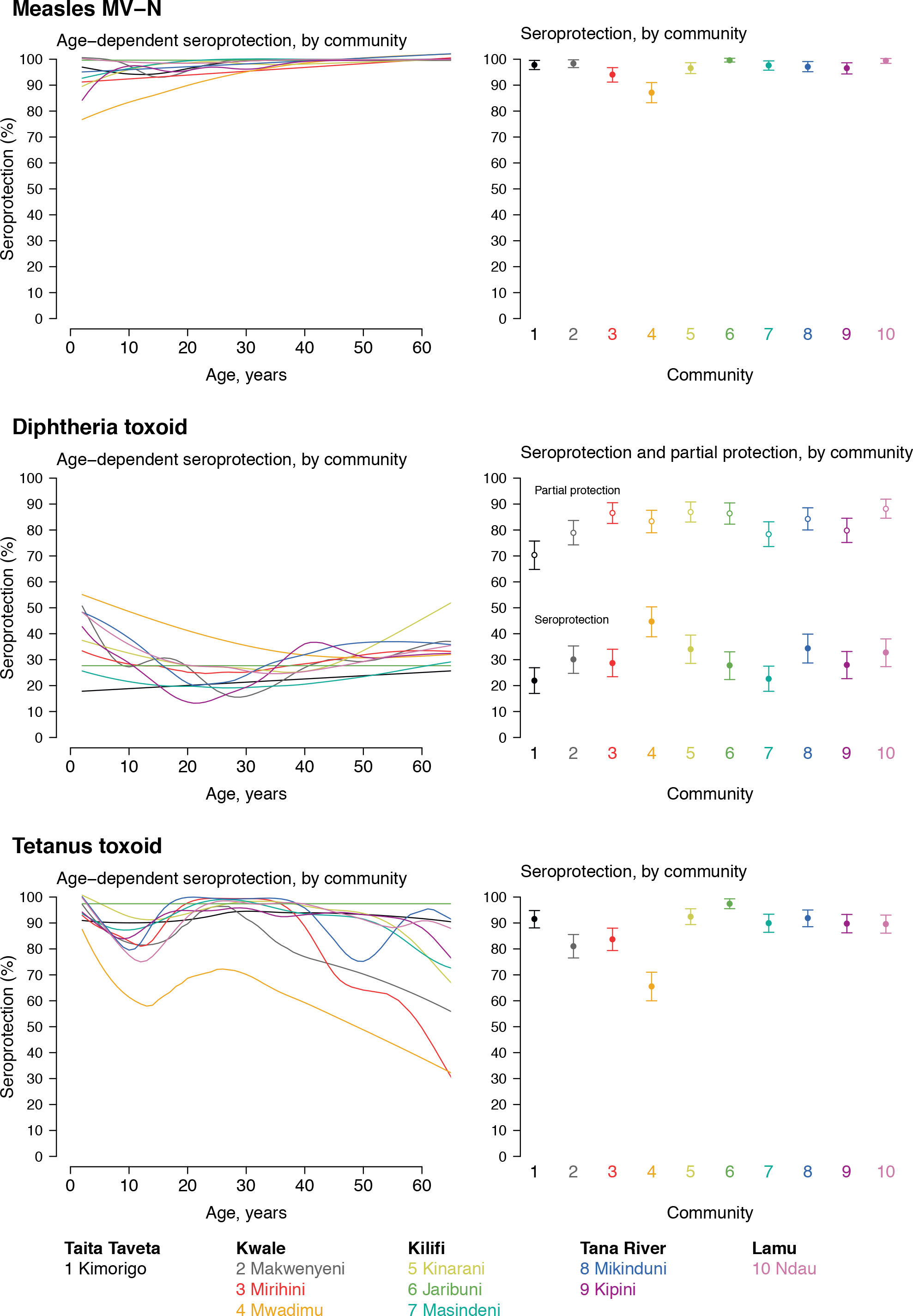
Age-dependent seroprotection and overall seroprotection for measles, diphtheria, and tetanus stratified by community in Kenya’s coastal region, 2015. Community-level seroprotection is age-adjusted and error bars represent 95% confidence intervals. For diphtheria, we included separate community level estimates of seroprotection (MFI > 4393 corresponding to 0.1 IU/ml) and partial protection (MFI > 183 corresponding to 0.01 IU/ml). Figure S6 is an extended version of this figure that also includes quantitative antibody levels. The script that created this figure is here: https://osf.io/qrkhm.

## Discussion

Many established national parasitic disease control and elimination programmes conduct routine surveillance to monitor and evaluate the impact of their targeted interventions. Epidemiologic surveillance systems that enable integrated surveillance and monitoring of co-endemic diseases and public health interventions could provide cost-effective synergy to support public health programs [20]. Antibodies can provide valuable information about past exposure to pathogens, and can be helpful for characterizing transmission dynamics in an area to help prioritize where and what interventions are needed the most [23,44].

The LF survey in coastal region of Kenya, which provided the opportunity to collect information for this study, demonstrated that Ndau Island in Lamu County had the highest prevalence of CFA by ICT [17]. The antifilarial antibody measurements assessed by MBAs closely aligned with the CFA results. Ndau Island had the highest levels of antibody responses to all three recombinant antifilarial antigens, which confirms the observation that LF transmission is currently higher in Ndau Island compared to the other communities. Previous studies have demonstrated a spatial relationship between antibody-positive individuals and infected persons [45]. The high seroprevalence rates in Ndau, especially among children, are consistent with the conclusion that transmission is ongoing and not yet halted by the MDA campaign.

Antibody responses against Bm14 antigen continued to increase with age in all villages, which may have been an indication of cumulative exposure to *W. bancrofti*, and also likely reflects historic transmission. Generally, antibody responses against the three recombinant filarial antigens were higher among with CFA-positive individuals than in CFA-negative persons although the difference was relatively smaller for Bm33 (see Fig. S7). Results from a recent study in American Samoa demonstrated that PCR-positive pools of LF vector mosquitoes were statistically significant predictors of seropositivity for Wb123 but not Bm14, suggesting Wb123 could be an indicator of ongoing transmission [46]. Longitudinal studies in areas of intense LF transmission have shown that children acquire infections early in life [47,48]. Additionally, previous studies have demonstrated that antibody response against infective stage filarial larvae antigen Wb123 is a specific measure of *Wuchereria bancrofti* infection, and reduction in both antibody prevalence and transmission is seen most clearly in young children [49,50]. Quantitative antifilarial antibody responses among youngest children (2-5 yr, 6-10 yr) provided much higher resolution distinctions between communities compared with seroprevalence using the same antigens or the ICT test (S8 Fig) – a result consistent with a recent analysis across diverse pathogens in low transmission settings where seropositive individuals are rare [43]. The higher resolution of quantitative antibody responses compared with seroprevalence, particularly when measured in small sampling clusters, suggests that quantitative antibody levels could serve as an important and more sensitive indicator of recent exposure in sentinel populations of young children, and may be valuable tool for surveillance in the context of lymphatic filariasis elimination programs [19]. Thus, combined measurement of these markers may be suitable for characterization of LF transmission settings particularly towards the end of the program when the infection prevalence is very low.

There was high heterogeneity in malaria seroprevalence among the study communities with Kwale and Kilifi counties generally showing relatively higher malaria transmission compared to the other 3 counties. The community mean seroprevalence values suggested that both *P. falciparum* and *P. malariae* transmission were highest in Jaribuni community in Kilifi County. These differences may reflect environmental heterogeneity in malaria larval breeding sites. A previous study in Kilifi and Kwale counties identified the primary vectors of malaria along the coast of Kenya to include *Anopheles funestus* and three members of the *An. gambiae* complex: *An. gambiae s.s.*, *An. arabiensis*, and *An. merus* [51]. The study also showed that relatively high malaria parasite prevalence can occur at low and even non-detectable levels of entomological inoculation rates (EIR), suggesting that measurement of EIR may be a relatively insensitive indicator of malaria transmission in some settings. Although malaria parasite prevalence and/or EIR have traditionally been used for reporting malaria transmission intensity [52], serological markers have increasingly been recognized as useful indicators for estimating malaria transmission intensity, which is key for assessing the impact of control interventions [53–56]. Because of the longevity of the specific antibody response, seroprevalence reflects cumulative exposure and thus is less affected by seasonality or unstable transmission [57].

In Kenya, *Schistosoma haematobium* is highly endemic along the coast where human exposure occurs primarily at pond and stream snail habitats [9,58,59]. The absence of *S. mansoni* from most of the Kenyan coastal region is attributable to the absence of the *Biomphalaria* spp. intermediate-host snails [60]. In Mikinduni Community, along the lower Tana River, crude antigen SEA antibody responses were observed, but *S. mansoni*-specific Sm25 responses were lacking. In contrast, Taveta area in Taita-Taveta County is known to be endemic for both *S. haematobium* and *S. mansoni* infections [61,62], and this is reflected in the high SEA and Sm25 antibody responses we observed in Kimorigo, a community located on the banks of the shallow freshwater Lake Jipe. The absence of *S. mansoni* species-specific antibody responses to Sm25 recombinant antigen in all of the communities except Kimorigo confirms that *S. mansoni* infection is likely absent from the lower coastal areas. Thus, *S. mansoni* Sm25 recombinant antigen seems to be an excellent antigen for measuring antibody responses to *S. mansoni* infection [63], and SEA antigen likely detects antibody responses caused by both *Schistosoma* species by virtue of cross-reactivity.

Presence of responses to *S. stercoralis* NIE antigen is noteworthy because there has been little information on the geographic distribution of this helminth in Kenya due to diagnostic limitations. Copromicroscopic diagnostic methods commonly used in soil-transmitted helminthiasis prevalence studies are inadequate for *S. stercoralis* detection [64], and thus its distribution in many areas is unknown. Concentration methods, namely the Baermann technique and Koga agar plate culture, have better but still unsatisfactory sensitivity [65]. A study employing NIE serology in Argentina found no cross-reactivity between *S. stercoralis* and infections with *A. lumbricoides*, hookworms, or *H. nana*, and the presence of other helminths in the stool did not affect the *S. stercoralis*-specific antibody responses [66]. A study comparing five serologic tests identified NIE-Luciferase Immunoprecipitation System to be the most accurate assay for the diagnosis of *S. stercoralis* infection [67]. Previous studies using the recombinant NIE have documented high seroprevalence of *S. stercoralis* infection in remote Australian Indigenous communities and suggest that collection of dried blood spots may be a useful approach for field diagnosis of *S. stercoralis* seroprevalence [68,69]. This study, therefore, provides evidence for possible low-level transmission of *S. stercoralis* in coastal Kenya as the seroprevalence varies from community to community. Community mean antibody responses to the *Ascaris* Hb native antigen and seroprevalence exhibited high heterogeneity among the study communities. A population-based study in Indonesia has shown that an assay for antibodies to *Ascaris* Hb is useful for assessing transmission of *Ascaris* infections, and community antibody rates decreased rapidly following MDA of anthelmintic drugs. The decrease was also found to reflect reduced egg excretion at the community level [33].

Vaccination is one of the most one of the most cost-effective public health interventions available, and the epidemiology and burden of vaccine-preventable diseases vary by country and by region partly because of differences in vaccine uptake [70]. This multiplex integrated serosurveillance study identified heterogeneity in serologic antibody levels against measles, diphtheria, and tetanus antigens. Our study demonstrates a need for regularly monitoring serological responses to vaccination programs in resource-poor settings where coverage may be low.

Some of the limitations of this study are somewhat similar to those highlighted previously [23]. Serological studies are traditionally faced with the challenge of establishing diagnostic cutoff points especially when well-characterized positive and negative serum samples are not available. Finite Gaussian mixture models applied in this study led to cutoff values that were very similar to those derived through ROC curves or from mean *plus* 3 standard deviation calculations for malaria, LF and helminth antibody responses (Fig 1). This result is consistent with a recent, multi-country comparison of cutoff methodology for trachoma antibodies [71], and supports the use of finite mixture models to identify seropositivity cutoffs in studies without access to panels of known positive and negative specimens. For pathogens where cutoff values fall in the centre of a unimodal distribution and it is more difficult to distinguish seropositive and seronegative groups (e.g., *A. suum* Hb in Fig 1), the use of community mean antibody levels avoids the requirement of choosing a cutoff, and observed antibody response patterns were very consistent with seroprevalence estimates across all of the antibodies tested in this study (S2–S6 Figs). Another limitation of this study is potential for antibody cross-reactivity. Since the coastal area has a typical tropical climate, it is likely that a plethora of pathogens are coincident, some with potentially cross-reactive antigens. A previous study reported that cross-reactivity of the *Ascaris* Hb native antigen with hookworm and possibly *S. stercoralis* and *Toxocara* spp. limited its value in serology if one is interested in ascariasis alone [33]. Thus, further studies are required to identify sensitive and specific recombinant antigens that could be used with more confidence in serological assays.

In spite of these limitations this study employed a single multiplex integrated serological assay and analysis methodology to measure antibody levels against several pathogens. There was no need to run separate assays for each pathogen, and we did not need to develop different mathematical models for each pathogen in order to compare exposure across communities and counties. The study highlighted overlap in pathogen burden that would not necessarily have been detected through single-disease surveillance. For example, Ndau Island was found to have the highest LF seroprevalence, but it also had highest *Ascaris* seroprevalence, thus supporting integrated control of these two helminths. Interestingly, Ndau had almost no evidence for *P. falciparum* malaria transmission. On the other hand Jaribuni community was found to stand out in terms of malaria, LF, and *Strongyloides*. Multiplex, integrated surveillance has the potential to enable us to look across diseases for opportunities for integrated control, thus providing synergy to global public health initiatives.

## Conclusion

This study highlighted the utility of the MBA platform for integrated serosurveillance of biomarkers of diseases of public health importance. The multiplex integrated serologic assay has the potential to become an invaluable tool for integrated monitoring of trends in endemicity of diseases of public health importance and the effectiveness of public health control programs.

## Acknowledgements

The authors would like to thank the County Health Departments of Taita-Taveta, Kwale, Kilifi, Tana River and Lamu for supporting the survey, including provision of laboratory technicians and local transportation for the survey teams. The communities of the selected sentinel sites and their local leaders are sincerely thanked for the cooperation and assistance. Drs. Patrick Lammie (CDC) and Simon Brooker (LSHTM) are thanked for useful comments and suggestions throughout the study. We wish to thank members of the Vaccine Preventable Disease Branch (CDC) including Sun Bae Sowers for sharing measles PRNT data. The Kenya Medical Research Institute (KEMRI) provided scientific leadership and oversight for this study.

## Disclaimer

Use of trade names is for identification only and does not imply endorsement by the Public Health Service or by the U.S. Department of Health and Human Services. The findings and conclusions in this report are those of the authors and do not necessarily represent the official position of the Centers for Disease Control and Prevention.

## Supporting Information

**S1 Checklist:** STROBE checklist.

STROBE Statement—Checklist of items that should be included in reports of ***cross-sectional studies***

|  | Item No | Recommendation |
| --- | --- | --- |
| **Title and abstract** | 1 | (a) Indicate the study's design with a commonly used term in the title or the abstract [The term cross-sectional is used within the title page 1, line 1 and the abstract page 2, line 29]<br>(b) Provide in the abstract an informative and balanced summary of what was done and what was found [page 2, lines 28-48] |
| **Introduction** |  |  |
| Background/rationale | 2 | Explain the scientific background and rationale for the investigation being reported [page 5, lines 96-110] |
| Objectives | 3 | State specific objectives, including any prespecified hypotheses [page 5, lines 106-110] |
| **Methods** |  |  |
| Study design | 4 | Present key elements of study design early in the paper [page 5, line 113] |
| Setting | 5 | Describe the setting, locations, and relevant dates, including periods of recruitment, exposure, follow-up, and data collection [page 5, lines 114-115] |
| Participants | 6 | (a) Give the eligibility criteria, and the sources and methods of selection of participants [page 5, lines 115-117] |
| Variables | 7 | Clearly define all outcomes, exposures, predictors, potential confounders, and effect modifiers. Give diagnostic criteria, if applicable [pages 8-9, lines 170-182] |
| Data sources/<br>measurement | 8* | For each variable of interest, give sources of data and details of methods of assessment (measurement). Describe comparability of assessment methods if there is more than one group [pages 8-9, lines 170-182] |
| Bias | 9 | Describe any efforts to address potential sources of bias [page 8, lines 184-217] |
| Study size | 10 | Explain how the study size was arrived at [page 5, lines 114-117] |
| Quantitative variables | 11 | Explain how quantitative variables were handled in the analyses. If applicable, describe which groupings were chosen and why [page 10, lines 219-228] |
| Statistical methods | 12 | (a) Describe all statistical methods, including those used to control for confounding [page 10, lines 223-225]<br>(b) Describe any methods used to examine subgroups and interactions [page 10, lines 217-219]<br>(c) Explain how missing data were addressed [N/A]<br>(d) If applicable, describe analytical methods taking account of sampling strategy [N/A]<br>(e) Describe any sensitivity analyses [N/A] |
| **Results** |  |  |
| Participants | 13* | (a) Report numbers of individuals at each stage of study—eg numbers potentially eligible, examined for eligibility, confirmed eligible, included in the study, completing follow-up, and analysed [page 10, lines 231-232; Figure S1]<br>(b) Give reasons for non-participation at each stage [N/A]<br>(c) Consider use of a flow diagram [N/A] |
| Descriptive data | 14* | (a) Give characteristics of study participants (eg demographic, clinical, social) and information on exposures and potential confounders [page 5, line 117]<br>(b) Indicate number of participants with missing data for each variable of interest [N/A] |
| Outcome data | 15* | Report numbers of outcome events or summary measures [pages 11-12; Figures |

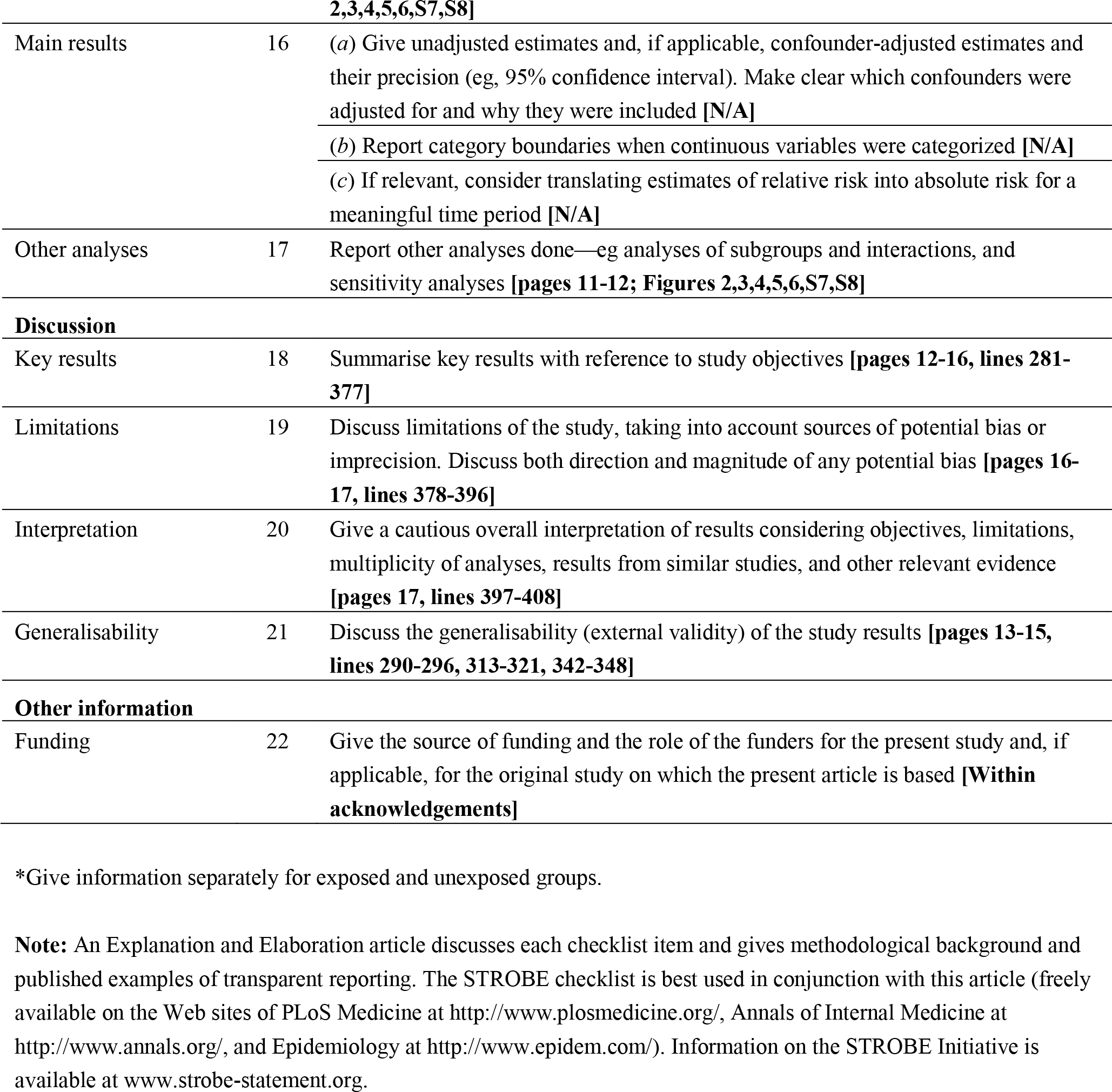

**2,3,4,5,6,S7,S8]**
|  |  |  |
| --- | --- | --- |
| Main results | 16 | (a) Give unadjusted estimates and, if applicable, confounder-adjusted estimates and their precision (eg, 95% confidence interval). Make clear which confounders were adjusted for and why they were included [N/A]<br>(b) Report category boundaries when continuous variables were categorized [N/A]<br>(c) If relevant, consider translating estimates of relative risk into absolute risk for a meaningful time period [N/A] |
| Other analyses | 17 | Report other analyses done—eg analyses of subgroups and interactions, and sensitivity analyses [pages 11-12; Figures 2,3,4,5,6,S7,S8] |
| **Discussion** |  |  |
| Key results | 18 | Summarise key results with reference to study objectives [pages 12-16, lines 281-377] |
| Limitations | 19 | Discuss limitations of the study, taking into account sources of potential bias or imprecision. Discuss both direction and magnitude of any potential bias [pages 16-17, lines 378-396] |
| Interpretation | 20 | Give a cautious overall interpretation of results considering objectives, limitations, multiplicity of analyses, results from similar studies, and other relevant evidence [pages 17, lines 397-408] |
| Generalisability | 21 | Discuss the generalisability (external validity) of the study results [pages 13-15, lines 290-296, 313-321, 342-348] |
| **Other information** |  |  |
| Funding | 22 | Give the source of funding and the role of the funders for the present study and, if applicable, for the original study on which the present article is based [Within acknowledgements] |

\*Give information separately for exposed and unexposed groups.
**Note:** An Explanation and Elaboration article discusses each checklist item and gives methodological background and published examples of transparent reporting. The STROBE checklist is best used in conjunction with this article (freely available on the Web sites of PLoS Medicine at http://www.plosmedicine.org/, Annals of Internal Medicine at http://www.annals.org/, and Epidemiology at http://www.epidem.com/). Information on the STROBE Initiative is available at www.strobe-statement.org.

**Table S1.** Seropositivity cutoff values estimated for each antigen. The cutoff values for vaccine preventable diseases (tetanus, diphtheria, and measles) are based on seroprotection levels derived from a standard curve using international reference. The cutoff values are plotted in Figure 1 of the manuscript.

| Antigen | Method | Cutoff (MFI-bg) |
| --- | --- | --- |
| Tetanus toxoid | Standard curve, Pos > 0.01 IU/ ml | 118 |
| Diphtheria toxoid | Standard curve, Pos > 0.1 IU/ ml | 4393 |
| Measles MV-N | Standard curve, Pos > 9.5 mIU/ml | 178 |
| *Plasmodium falciparum* CSP | Gaussian mixture model <sup>a</sup> | 367 |
| *Plasmodium falciparum* MSP-1<sub>19</sub> | USA unexp. pop., mean + 3SD of log values | 256 |
| *Plasmodium falciparum* MSP-1<sub>19</sub> | Gaussian mixture model <sup>a</sup> | 416 |
| *Plasmodium malariae* MSP-1<sub>19</sub> | USA unexp. pop., mean + 3SD of log values | 403 |
| *Plasmodium malariae* MSP-1<sub>19</sub> | Gaussian mixture model <sup>a</sup> | 196 |
| Lymphatic filariasis Wb123 | USA unexp. pop., mean + 3SD | 342 |
| Lymphatic filariasis Wb123 | Gaussian mixture model <sup>a</sup> | 366 |
| Lymphatic filariasis Bm14 | USA unexp. pop., mean + 3SD | 444 |
| Lymphatic filariasis Bm14 | Gaussian mixture model <sup>a</sup> | 214 |
| Lymphatic filariasis Bm33 | USA unexp. pop., mean + 3SD | 369 |
| Lymphatic filariasis Bm33 | Gaussian mixture model <sup>a</sup> | 791 |
| *Schistosoma mansoni* SEA | ROC analysis with pos/neg samples | 965 |
| *Schistosoma mansoni* SEA | Gaussian mixture model <sup>a</sup> | 476 |
| *Schistosoma mansoni* Sm25 | ROC analysis with pos/neg samples | 38 |
| *Schistosoma mansoni* Sm25 | Gaussian mixture model <sup>a</sup> | 32 |
| *Strongyloides stercoralis* NIE | USA unexp. pop., mean + 3SD | 628 |
| *Strongyloides stercoralis* NIE | Gaussian mixture model <sup>a</sup> | 645 |
| *Ascaris* AsHb | ROC analysis with pos/neg samples | 386 |
| *Ascaris* AsHb | Gaussian mixture model <sup>a</sup> | 780 |

<sup>a</sup> Model fit using the mixtools package in R on the original scale of the antibody measurement, and assuming a mixture of 2 components. The mean of the lower component + 3SD was used to determine cutoff values. The script used to run the mixture model analyses is here: https://osf.io/dm324, and the log output from the analysis (estimates copied/pasted to table) is here: https://osf.io/ms39q

**S1 Figure.**
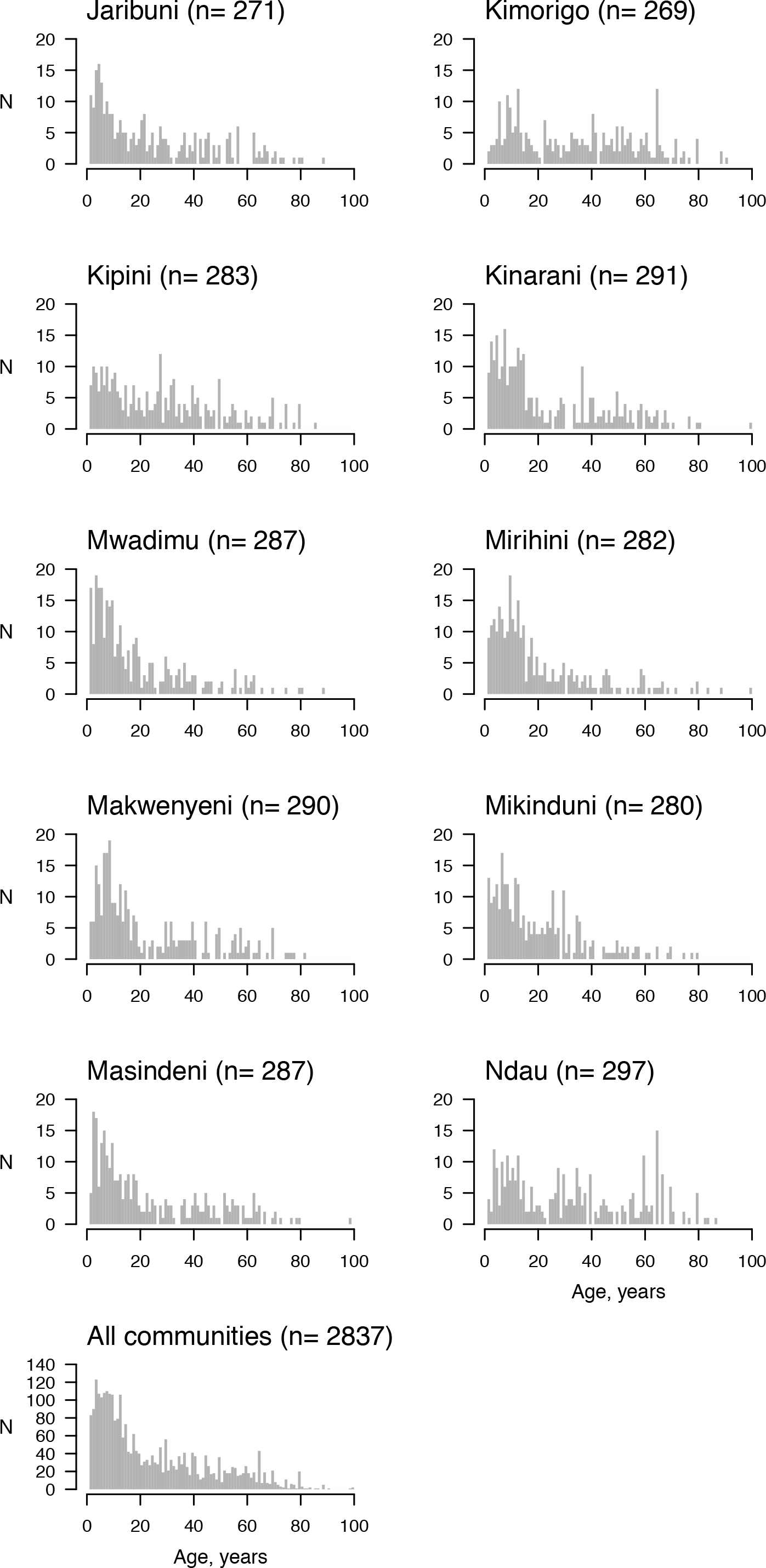
Community level sample size and age distribution. Figure created with computational notebook: https://osf.io/7jxmn.

**S2 Figure:**
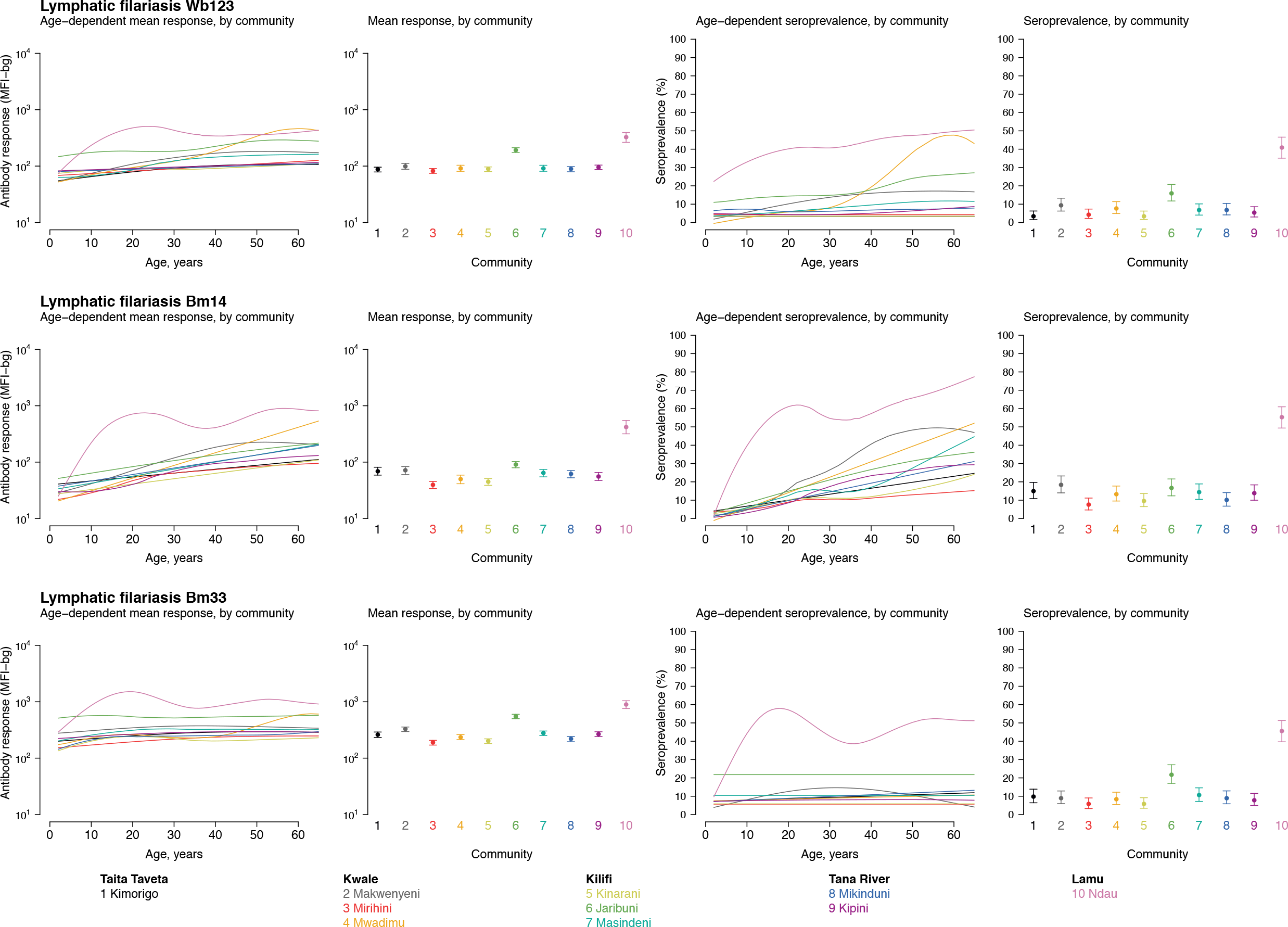
Lymphatic filariasis antibody age-dependent mean response and seroprevalence, stratified by community in Kenya?s coastal region, 2015. Community-level mean antibody response and seroprevalence are age-adjusted and error bars represent 95% confidence intervals. Antibody response measured in median fluorescence units minus background (MFI-bg) on a BioRad Bio-Plex platform. Figure created with computational notebook: https://osf.io/c79rw.

**S3 Figure:**
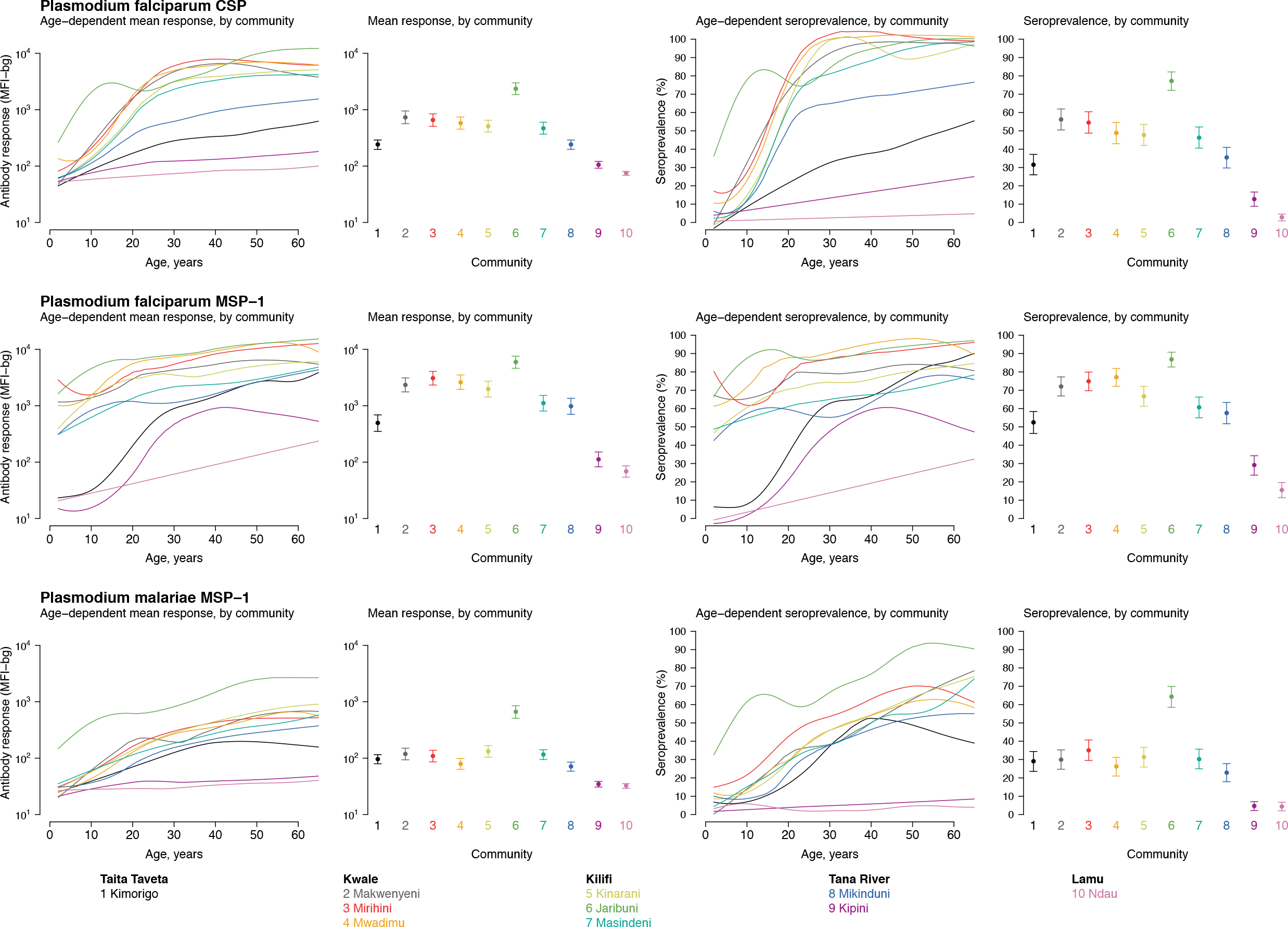
Malarial antibody age-dependent mean response and seroprevalence, stratified by community in Kenya?s coastal region, 2015. Community-level mean antibody response and seroprevalence are age-adjusted and error bars represent 95% confidence intervals. Antibody response measured in median fluorescence units minus background (MFI-bg) on a BioRad Bio-Plex platform. Figure created with computational notebook: https://osf.io/nhrc2.

**S4 Figure:**
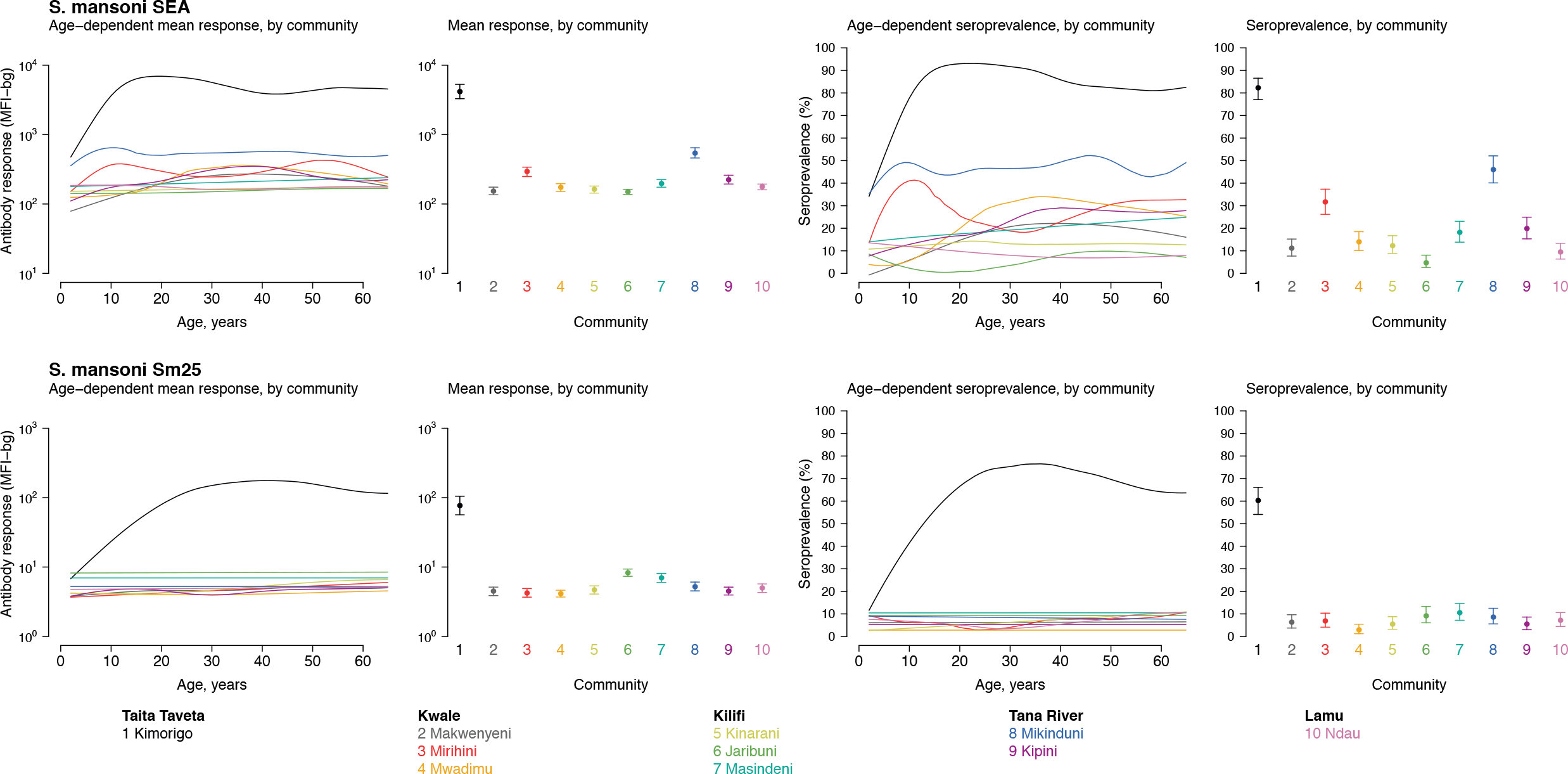
Schistosomiasis antibody age-dependent mean response and seroprevalence, stratified by community in Kenya?s coastal region, 2015. Community-level mean antibody response and seroprevalence are age-adjusted and error bars represent 95% confidence intervals. Antibody response measured in median fluorescence units minus background (MFI-bg) on a BioRad Bio-Plex platform. Figure created with computational notebook: https://osf.io/z8v4n.

**S5 Figure:**
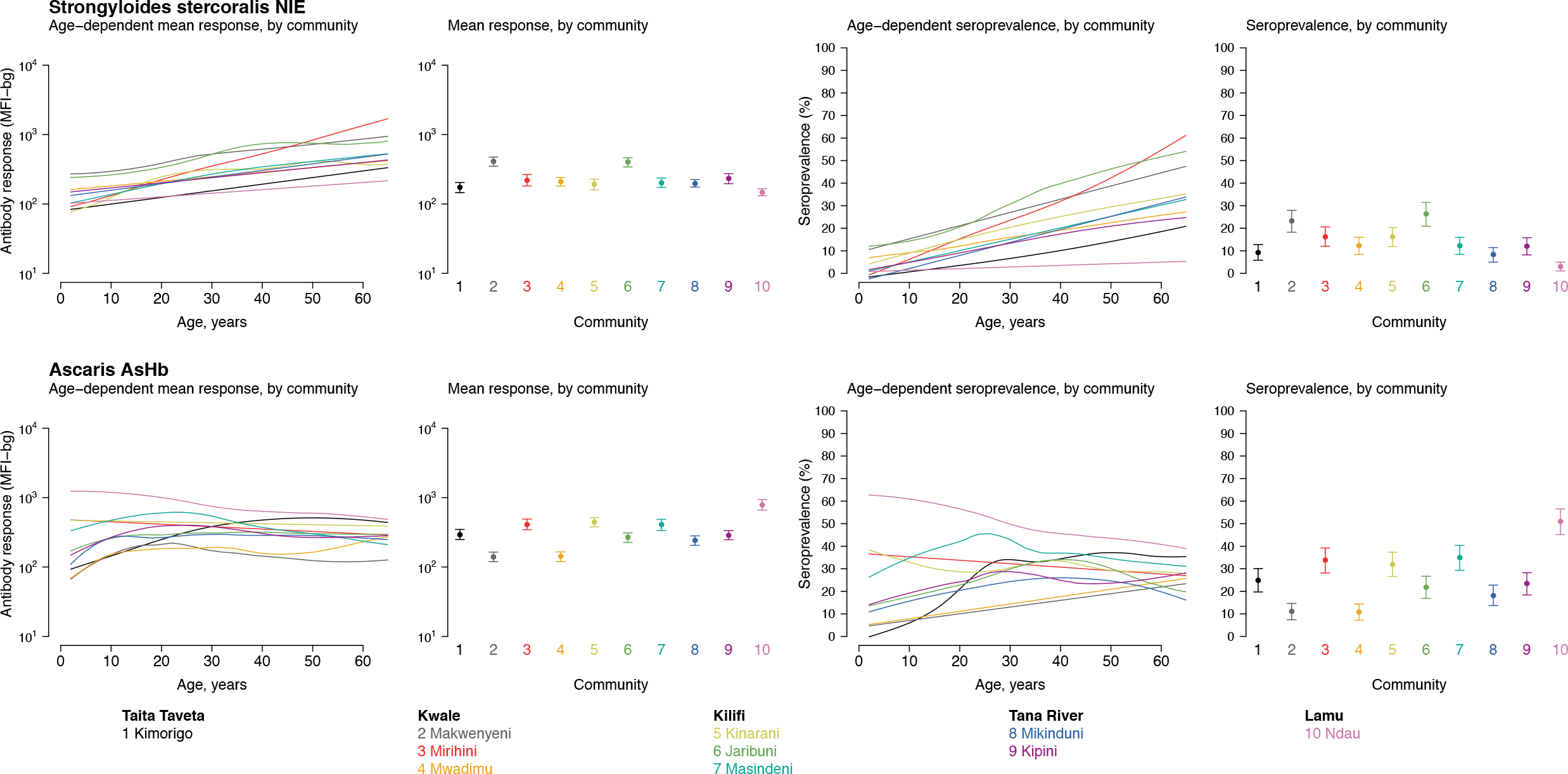
Age-dependent mean response and seroprevalence antibodies to S. stercoralis and A. lumbricoides, stratified by community in Kenya?s coastal region, 2015. Community-level mean antibody response and seroprevalence are age-adjusted and error bars represent 95% confidence intervals. Antibody response measured in median fluorescence units minus background (MFI-bg) on a BioRad Bio-Plex platform. Figure created with computational notebook: https://osf.io/spnvx.

**S6 Figure:**
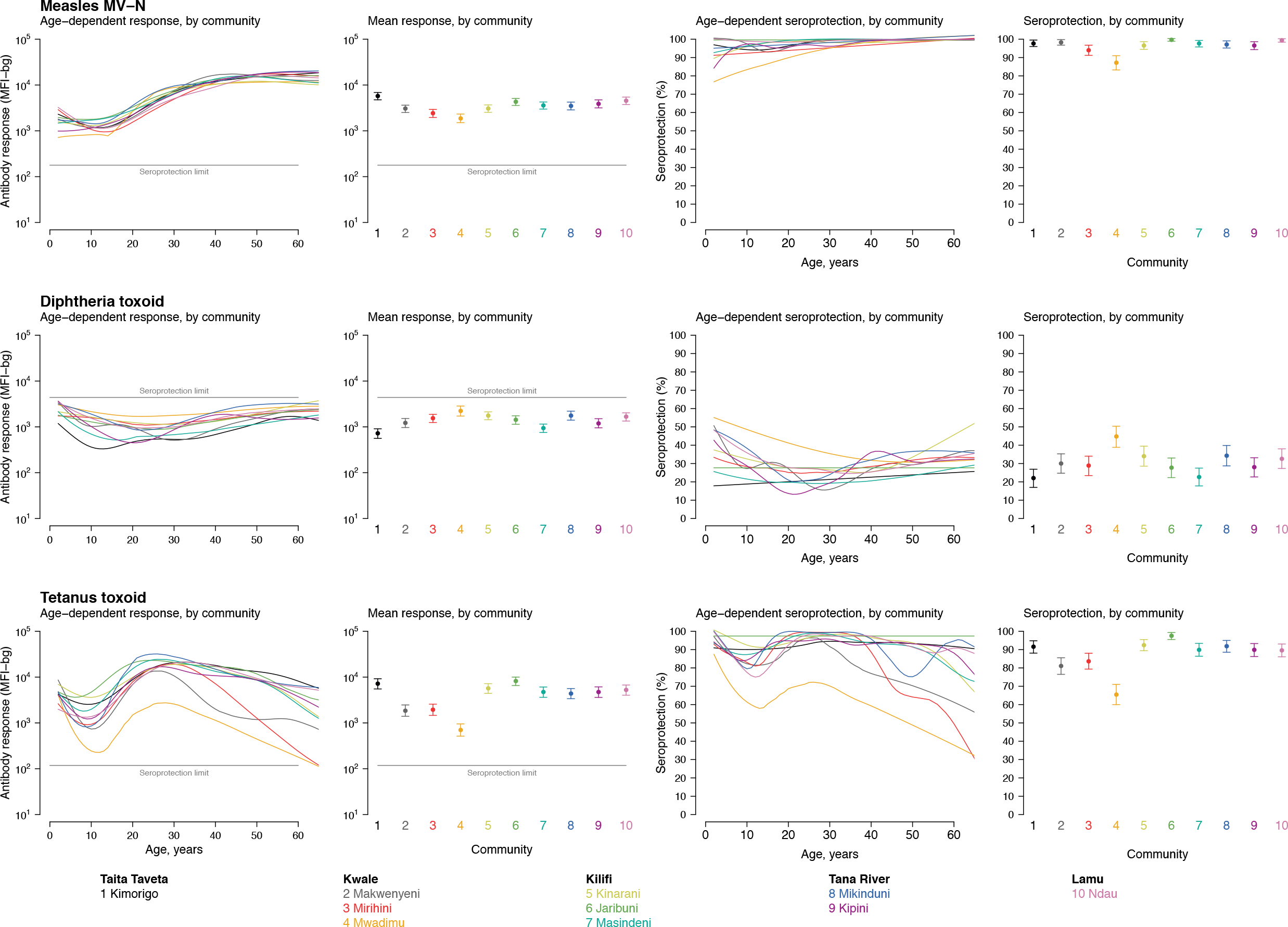
Age-dependent mean response and seroprotection for measles, diphtheria, and tetanus stratified by community in Kenya?s coastal region, 2015. Community-level mean antibody response and seroprotection are age-adjusted and error bars represent 95% confidence intervals. Antibody response measured in median fluorescence units minus background (MFI-bg) on a BioRad Bio-Plex platform. Figure created with computational notebook: https://osf.io/uy5bf.

**S7 Figure:**
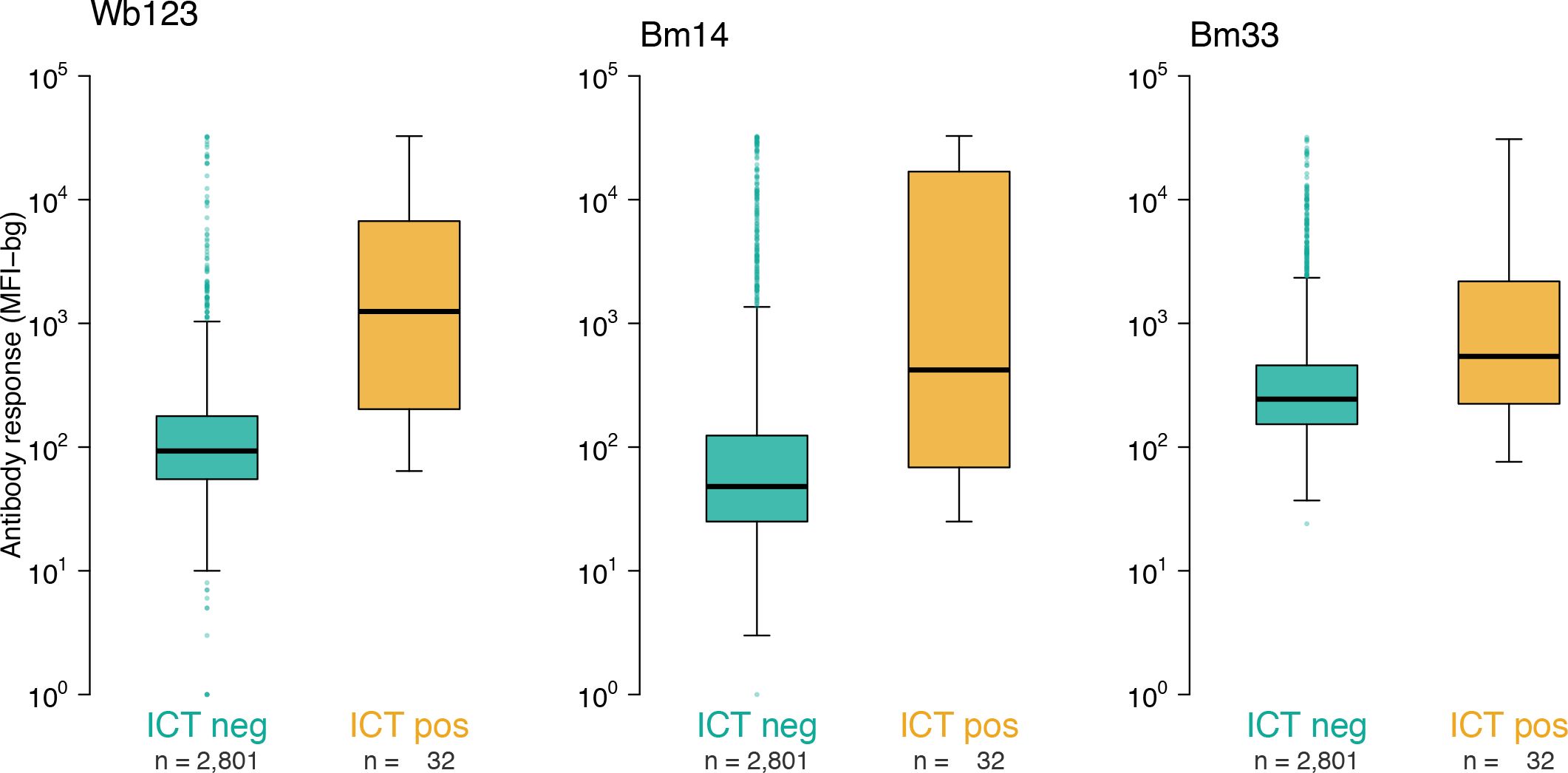
Distribution of three lymphatic filariasis antibodies, stratified by rapid antigen immunochromatographic card test (ICT) results. Boxes mark the median and interquartile range of the distributions. Antibody response measured in median fluorescence units minus background (MFI-bg) on a BioRad Bio-Plex platform. Mann-Whitney U-test P < 0:0001 for differences in antibody responses between ICT negative and positive individuals. Figure created with computational notebook: https://osf.io/k9tms.

**S8 Figure:**
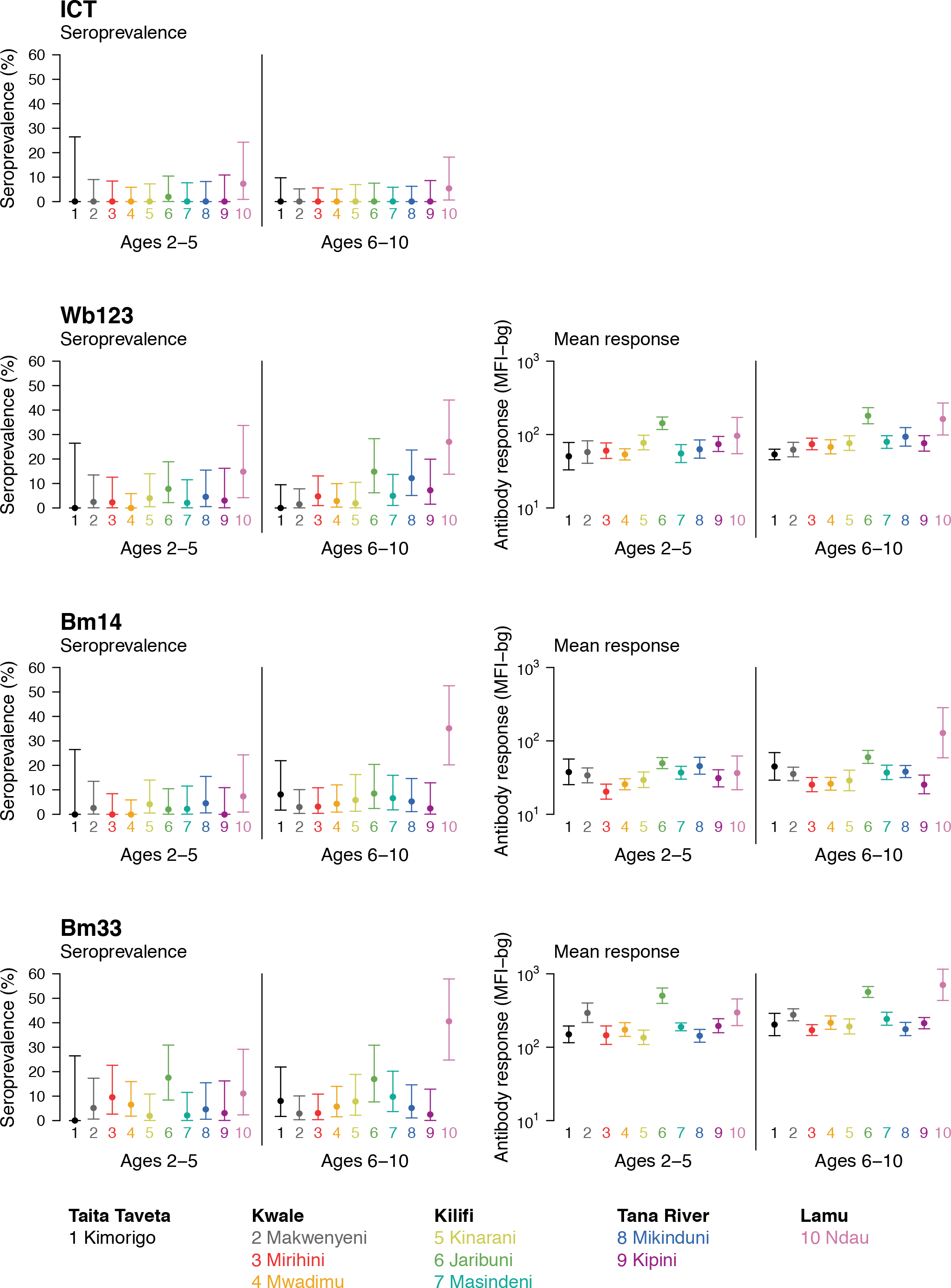
Community level estimates of lymphatic filariasis seroprevalence and geometric mean antibody levels among children ages 2-5 and 6-10 years old. Child blood samples were tested the immunochromatographic card test (ICT) and three antigens (Wb123, Bm14, Bm33) measured in median fluorescence units minus background (MFI-bg) on a multiplex BioRad Bio-Plex platform. The mean number of specimens tested per community within each age stratum was 47 (median=47; interquartile range= 39, 58; range= 12, 70). Figure created with computational notebook: https://osf.io/xh9yt.

